# A nucleus-like compartment shields bacteriophage DNA from CRISPR-Cas and restriction nucleases

**DOI:** 10.1101/370791

**Authors:** Senén D. Mendoza, Joel D. Berry, Eliza S. Nieweglowska, Lina M. Leon, David A. Agard, Joseph Bondy-Denomy

## Abstract

**All viruses require strategies to inhibit or evade the immunity pathways of cells they infect. The viruses that infect bacteria, bacteriophages (phages), must avoid nucleic-acid targeting immune pathways such as CRISPR-Cas and restriction endonucleases to replicate efficiently^1^. Here, we show that a jumbo phage infecting *Pseudomonas aeruginosa*, phage ΦKZ, is resistant to many immune systems *in vivo*, including CRISPR-Cas3 (Type I-C), Cas9 (Type II-A), Cas12 (Cpf1, Type V-A), and Type I restriction-modification (R-M) systems. We propose that ΦKZ utilizes a nucleus-like shell to protect its DNA from attack. Supporting this, we demonstrate that Cas9 is able to cleave ΦKZ DNA *in vitro*, but not *in vivo* and that Cas9 is physically occluded from the shell assembled by the phage during infection. Moreover, we demonstrate that the Achilles heel for this phage is the mRNA, as translation occurs outside of the shell, rendering the phage sensitive to the RNA targeting CRISPR-Cas enzyme, Cas13a (C2c2, Type VI-A). Collectively, we propose that the nucleus-like shell assembled by jumbo phages enables potent, broad spectrum evasion of DNA-targeting nucleases.**

---

Phage infection and replication can cause bacterial death via cell lysis, necessitating protective immune systems to negate this great threat to bacterial viability^2,3^. Restriction-modification (R-M) and adaptive CRISPR-Cas (clustered regularly interspaced short palindromic repeats and CRISPR-associated genes) immunity detect and degrade phage nucleic acids to protect the host^1^. Recent CRISPR-Cas discovery efforts have led to the characterization of six distinct CRISPR-Cas types (I-VI) with independent mechanisms for CRISPR RNA (crRNA) biogenesis, surveillance complex assembly, substrate selection, and target degradation^4^. Type III and VI CRISPR-Cas systems target RNA^5,6^, while Types I, II, and V predominantly target DNA^7-9^. Despite these differences, all characterized CRISPR-Cas systems function via crRNA-guides derived from the DNA-based CRISPR array, which stores the memory of past infections^10,11^.

Phages that infect *Pseudomonas aeruginosa* avoid CRISPR-mediated destruction by encoding “anti-CRISPR" (Acr) proteins that inhibit the Type I-E and I-F CRISPR-Cas systems^12-14^. We sought to determine whether any *P. aeruginosa* phages are also resistant to the Type I-C CRISPR-Cas subtype, also present in *P. aeruginosa*^15^. This subtype is notable as it is the most minimal Type I system identified to date, is highly abundant in bacterial genomes^16^, and is understudied relative to other subtypes. We identified an isolate in our lab encoding a Type I-C system, and designed and expressed a crRNA targeting related phages JBD30 and DMS3m, which encode AcrIF and AcrIE proteins, respectively. The successful crRNA-specific targeting of these phages by the Type I-C system demonstrated functional CRISPR immunity and that these phages did not possess functional Type I-C Acr proteins (Fig. 1a). To test the sensitivity of other phage families against the Type I-C system, we transferred the necessary *cas* genes (*cas3, cas5, cas8, cas7*) into the chromosome of a commonly used, phage-sensitive lab strain (PAO1) that naturally lacks CRISPR-Cas immunity. A small screen was then conducted, where crRNAs were tested against phages from five taxonomic groups: JBD30, D3, ΦKZ, F8, and JBD68. JBD30, D3, and JBD68 are distinct temperate siphophages while ΦKZ and F8 are distinct lytic myophages. All phages succumbed to targeting, except ΦKZ (Fig. 1b, 1c). ΦKZ titer did not decrease when exposed to ten different Type I-C crRNAs (Fig. 1c, Extended Data Fig. 1), suggesting that it is completely resistant to this immune system of *P. aeruginosa*.

**Figure 1:**
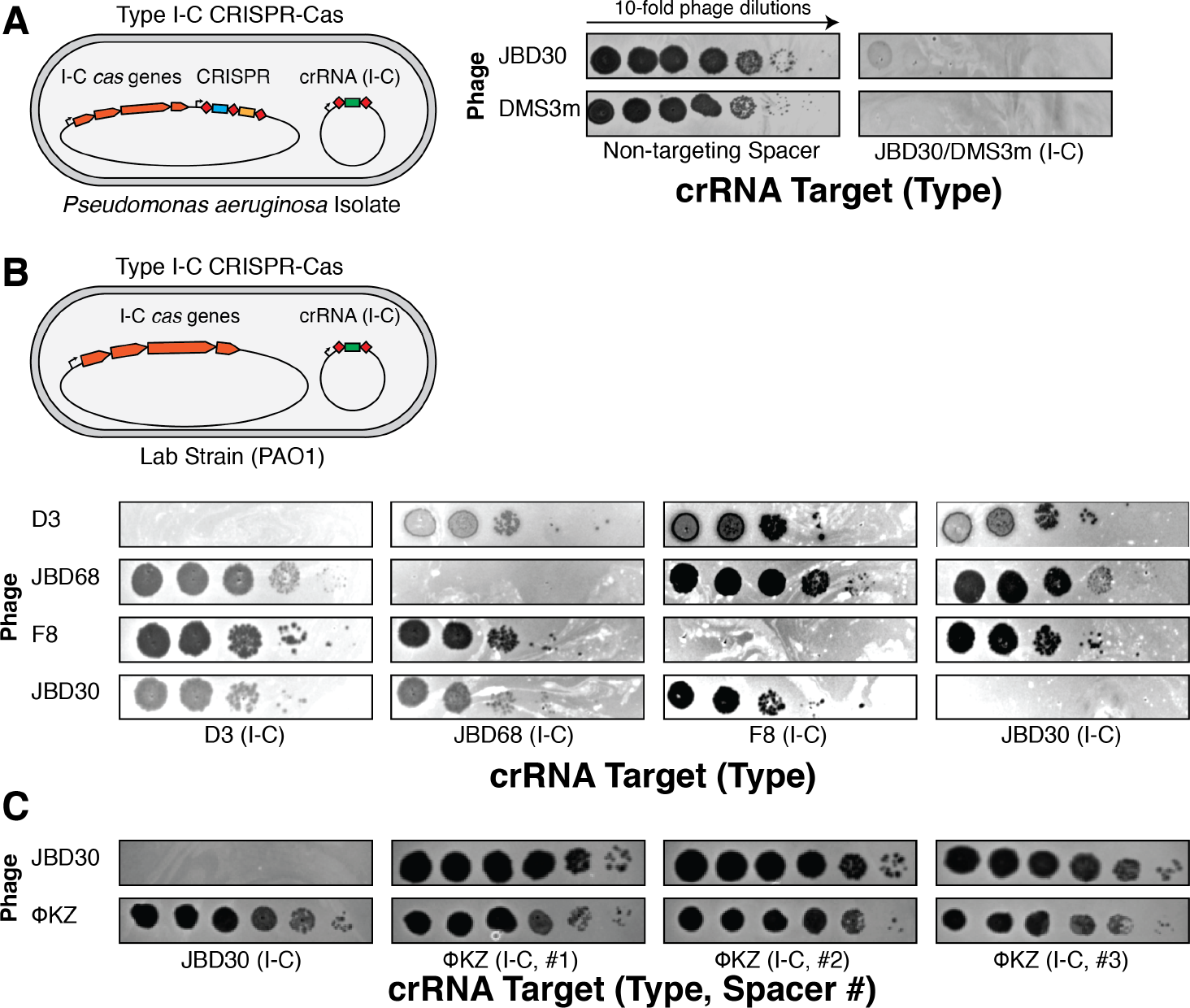
Identification of a phage that resists *P. aeruginosa* Type I-C CRISPR-Cas immunity. **a**, Phages JBD30 and DMS3m were spotted in ten-fold serial dilutions on a lawn of a *P. aeruginosa* isolate naturally expressing the I-C *cas* genes, and an engineered crRNA to target both phages. Dark clearings in the lawn represent phage replication. **b,** Strain PAO1 was engineered to express the I-C *cas* genes and crRNAs targeting the indicated phages, and plaque assays were conducted as in Fig. 1a. **c,** PAO1 strains expressing crRNAs engineered to target phage JBD30 and phage ΦKZ (I-C, #1-#3) were subjected to a plaque assay as in Fig. 1a.

The ΦKZ genome possesses no homologs of *acr* genes^12-14,17,18^ or anti-CRISPR associated (*aca*) genes that have previously enabled identification of *acr* genes^14,17,18^. Moreover, ΦKZ is a phage with no genetic tools currently available to manipulate it, rendering traditional knockout approaches infeasible. Thus to determine the mechanism by which the ΦKZ phage resists Type I CRISPR-Cas, we attempted to utilize the Type II-A CRISPR-Cas9 system from *Streptococcus pyogenes* (SpyCas9) to knock out phage genes. SpyCas9 and sgRNAs were adapted for expression and function in *P. aeruginosa,* leading to robust targeting of control phage JBD30, but notably, ΦKZ replication and associated cell lysis was unaffected both in plate and liquid assays (Fig. 2a). An additional eight sgRNA sequences also failed to target ΦKZ (Extended Data Fig. 2a), suggesting that ΦKZ resists both Type I and II CRISPR immunity. Given the ability of this phage to evade two unrelated CRISPR systems originating from distantly related microbes, we considered that it may be generally resistant CRISPR-Cas immunity. To test this, the Type V-A Cas12a (Cpf1) CRISPR-Cas system from *Moraxella bovoculi* was expressed in *P. aeruginosa* and again, robust targeting of phage JBD30 was observed, but not of ΦKZ with any of the nine crRNAs tested (Fig. 2b, Extended Data Fig. 2b). The ability of this phage to resist CRISPR systems found in its natural host (Type I-C) and those not present in *Pseudomonas* (Type II-A and V-A) suggests that this phage possesses a mechanism to enable “pan-CRISPR” resistance.

**Figure 2:**
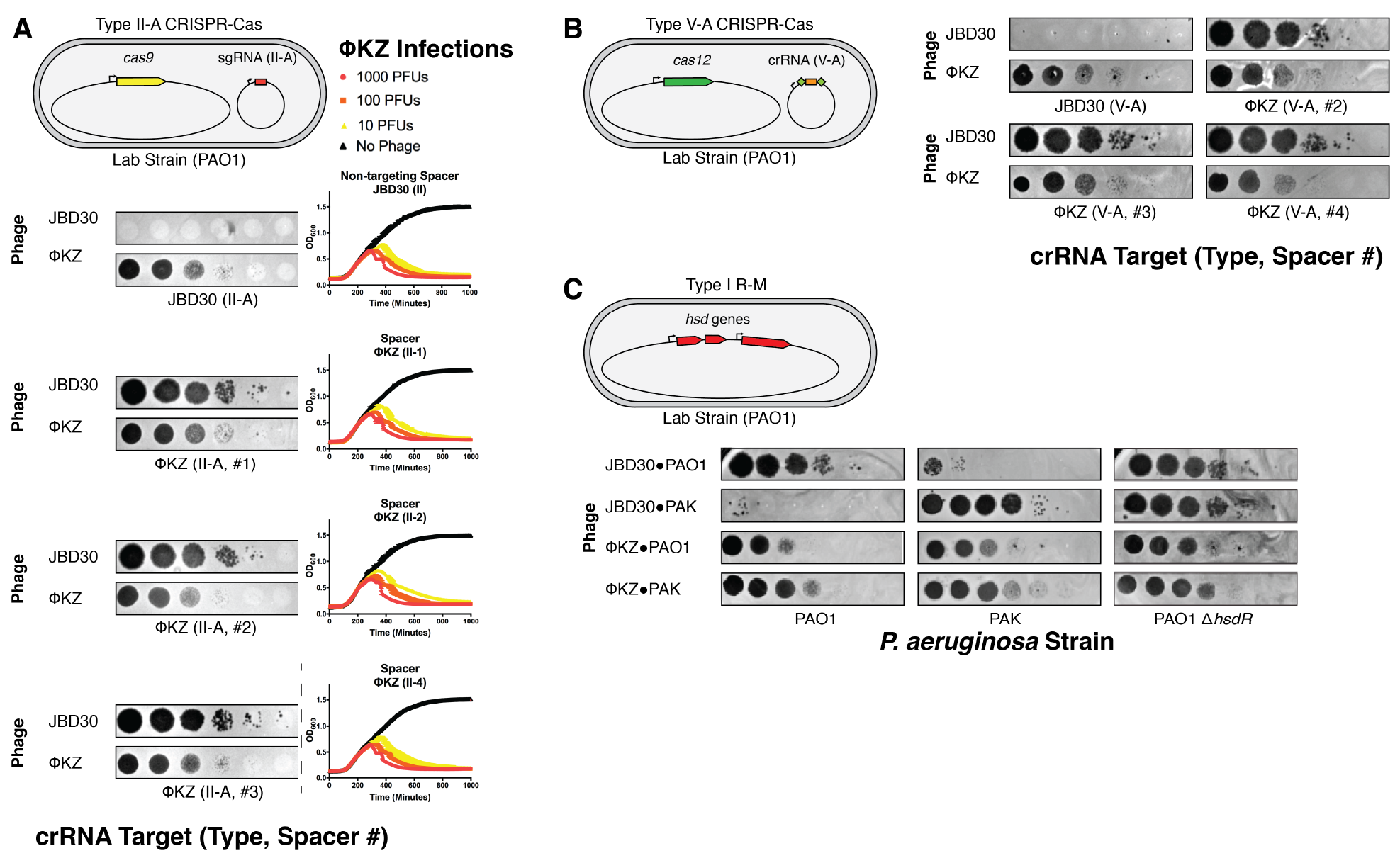
Jumbo phage ΦKZ resists targeting by heterologous Type II-A and V-A CRISPR-Cas systems and and endogenous R-M system. **a**, Strain PAO1 was engineered to express the Type II-A Cas9 protein and single guide RNAs (sgRNAs) targeting the indicated phage. Plaque assays were conducted as in Figure 1a and growth curves were conducted, monitoring the OD600 of PAO1 cells infected with the indicated number of ΦKZ plaque forming units (pfu). **b,** Strain PAO1 was engineered to express the Type V-A Cas12a protein and crRNAs against the indicated phage. Plaque assays conducted as in Figure 1a. **c,** The endogenous Type I R-M system (*hsdRSM*) in strains PAO1 and PAK was assayed using phages propoagated on PAO1 or PAK as indicated (e.g. JBD30•PAO1 was first propagated on strain PAO1). Together with an isogenic PAO1*∆hsdR* knockout, all strains were subjected to a plaque assay as in Figure 1a.

Restriction-modification systems are the most common bacterial immune system in nature and pose a significant impediment to phage replication^1^. To test whether ΦKZ is also resistant to attack from restriction endonucleases, the phage was propagated on strain PAK, an isolate that generates phages that are restricted by strain PAO1. When phage JBD30 was propagated on PAK, and then plated on strain PAO1, its titer was reduced by ~3 orders of magnitude (Fig. 2c), an effect that was ameliorated in a PAO1 strain lacking the Type I R-M system (*∆hsdR*). However, when ΦKZ was propagated on PAK, its titer did not decrease on PAO1 (Fig. 2c). Conversely, when propagated on PAO1, ΦKZ titer was not decreased on PAK, while JBD30 was again reduced in titer by ~3 orders of magnitude (Fig. 2c). This demonstrates that ΦKZ is also recalcitrant to Type I restriction endonucleases *in vivo*.

Given the strong resistance to targeting, we considered whether DNA base modifications were protecting this phage from enzyme targeting, as has been previously seen with phage T4 and others^19-22^. Purified phage DNA, extracted from ΦKZ virions, was subjected to restriction digestion reactions with a panel of restriction enzymes that are inhibited by glc-HmC moieties, including HindIII, EcoRI, SacI, KpnI, NcoI, and EcoRI (ref. 22,23 and per New England Biolabs). Additionally, MspJI was utilized, a modification-dependent restriction nuclease that requires 5-HmC or 5-mC modified DNA for cleavage^24^. Except for SacI, which lacks a sequence recognition motif in the ΦKZ genome, all enzymes tested cleaved ΦKZ gDNA, demonstrating the absence of glc-HmC modifications (Fig. 3a). To determine whether the phage genome is a substrate for CRISPR-Cas9-based cleavage, purified phage DNA was subjected to a cleavage assay with two distinct crRNA sequences using a dual crRNA:tracrRNA-loaded SpyCas9 nuclease *in vitro*. Cas9 cleaved the phage genome in two locations, liberating the predicted 10 kb fragment from the much larger 280 kb genome (Fig. 3b). Notably, the two crRNA sequences used here are the same sequence as crRNAs II-A, #1 and II-A, #2 used *in vivo* (Fig. 2a), demonstrating that these crRNA sequences do not target phage *in vivo,* but are competent for genome cleavage *in vitro.*

**Figure 3:**
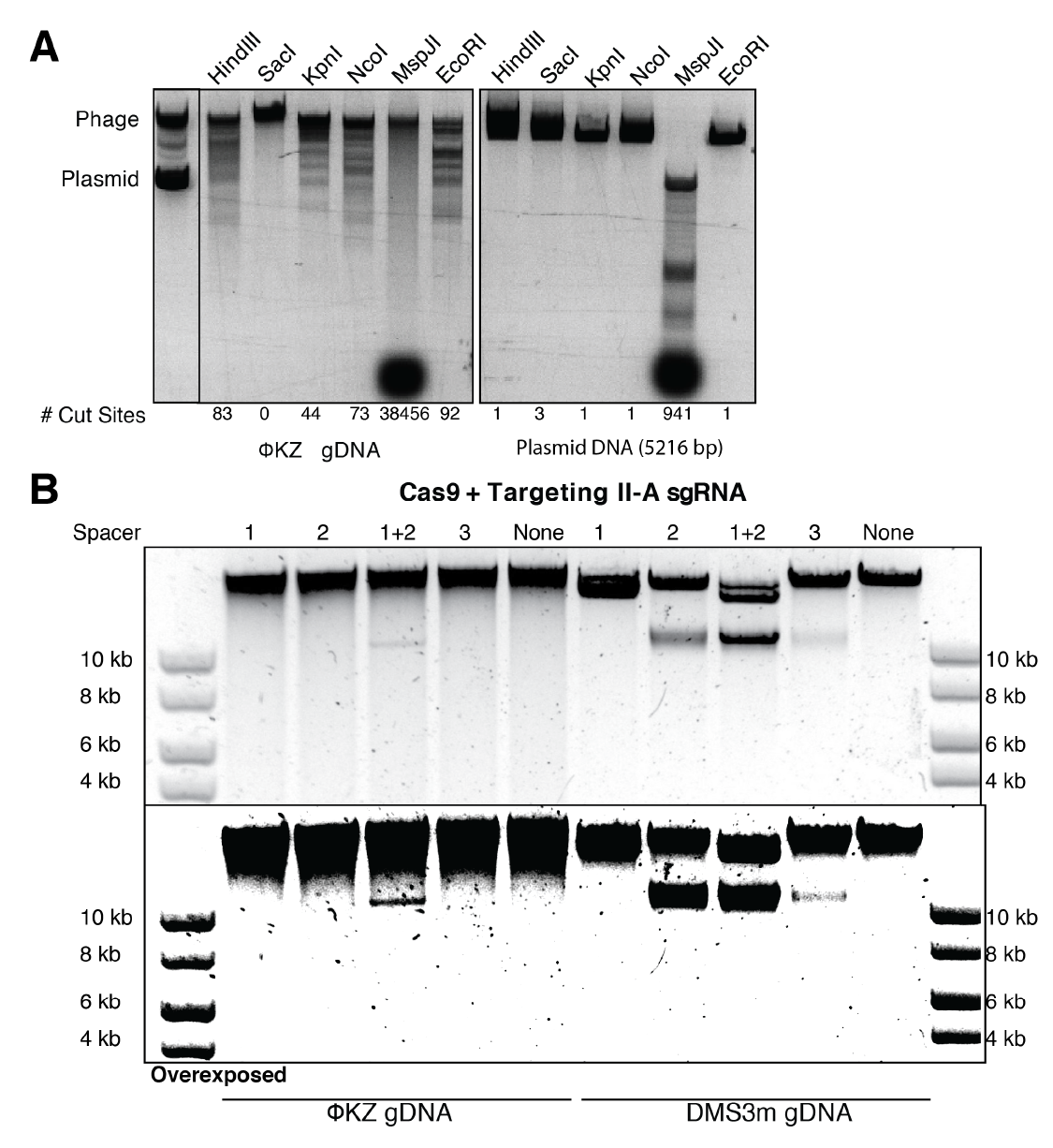
ΦKZ genomic DNA is sensitive to restriction enzymes and Cas9 *in vitro.* **a**, ΦKZ genomic DNA and plasmid DNA were subjected to digestion with the indicated restriction enzymes *in vitro*. The first lane contains purified phage and plasmid DNA run together. The number of cut sites for each enzyme is shown at the bottom of the gels. **b,** ΦKZ and phage DMS3m genomic DNA were digested *in vitro* using Cas9 loaded with crRNA:tracrRNA targeting the indicated phage. The bottom gel is the same as the top gel, however the image was overexposed to enhance faint bands. Products were visualized on a 0.7% agarose gel, visualized with SYBR Safe nucleic acid stain.

Recently it was shown that ΦKZ and ΦKZ-like phages infecting *Pseudomonas* sp. construct an elaborate nucleus-like, proteinaceous compartment where phage DNA replicates, with PhuZ, a phage tubulin homologue, centering the compartment within the host cell^25-29^. Proteins involved in DNA replication and transcription localize inside the shell, while proteins mediating translation and nucleotide synthesis are relegated to the cytoplasmic space, akin to the eukaryotic nucleus. Given the rapid assembly of the shell and its apparent exclusion of select proteins, we considered whether this structure was responsible for the resistance of ΦKZ to four unrelated CRISPR-Cas and restriction nucleases. The Type II-A single protein effector Cas9 was chosen as a representative immune system for ease of manipulation and imaging. *P. aeruginosa* cells infected with ΦKZ were imaged with a Cas9 fluorescent antibody, revealing that the protein is indeed excluded from the shell during phage infection (Fig. 4a). DAPI staining reveals the phage DNA inside the nucleus-like shell, while the host genome is rapidaly degraded during infection. As a control, a protein previously shown to be internalized in the shell, ORF152, was imaged revealing co-localization with the DAPI-positive phage nucleus. While the rules for protein internalization in the shell are currently unknown, these data and work from the initial shell studies^28,29^ suggest that the default localization for large host-encoded proteins is to be excluded from the shell.

**Figure 4:**
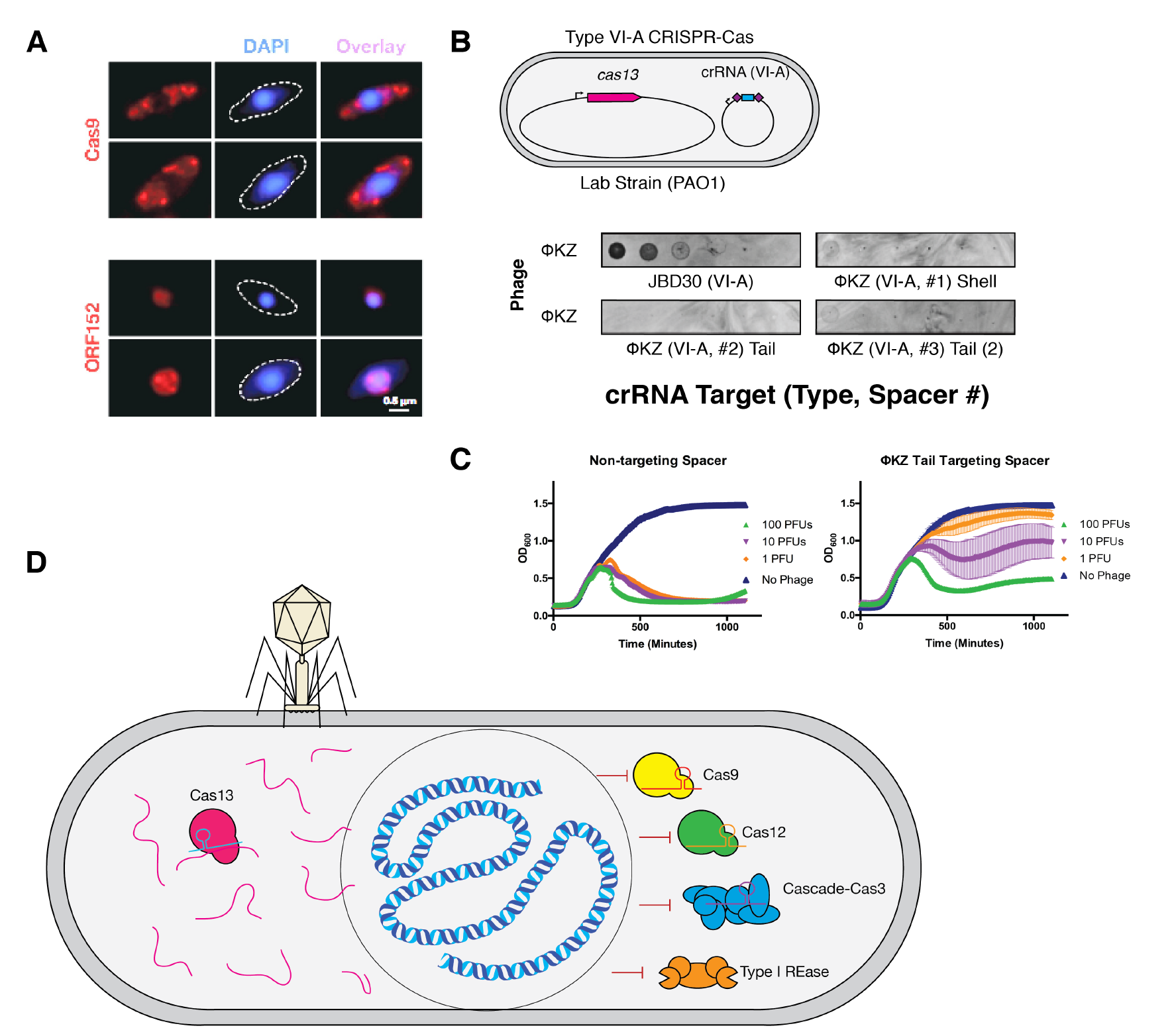
Phage ΦKZ DNA is protected from Cas9 and other DNA targeting enzymes, but is sensitive to RNA-targeting Cas13. **a**, Fluorescence microscopy of *P. aeruginosa*, immunostained for Cas9 (top panels), or Myc-ORF152 (bottom panels). DAPI stain shows the phage DNA within the shell. **b,** Strain PAO1 expressing LseCas13a and a crRNA targeting the indicated phage. Plaque assays conducted as in Figure 1a. **c,** Growth curves measuring the OD600 of PAO1 infected with the indicated number of ΦKZ plaque forming units (PFUs). **d,** A model summarizing the ΦKZ nucleus-like shell excluding Cas9, Cas12, Cascade-Cas3 (Type I-C, and Type I restriction endonucleases (REase), while the mRNA (red) is exported and can be targeted by Cas13.

While ΦKZ DNA is protected from CRISPR-Cas and restriction endonucleases, the mRNA is not afforded this same luxury, as it leaves the confines of the shell to be translated in the cytoplasm. We therefore envisaged that an RNA-targeting CRISPR-Cas system would provide immunity to ΦKZ even if localized in the cytoplasm. To test this, we adapted the RNA-guided RNA nuclease Cas13a (C2c2, Type VI-A) from *Listeria seeligeri*^5,30^ for phage targeting in *P. aeruginosa*. By targeting several different transcripts, three LseCas13 spacers were found (two targeting tail gene *gp146* and one targeting the shell gene *gp054*) that decreased ΦKZ plaquing efficiency by 10-1000-fold (Fig. 4b, Extended Data Fig. 3). Corroborating the plaquing results, LseCas13 also provided protection of *P. aeruginosa* in liquid cultures, with strong bacterial growth at phage inputs that kill cultures with a non-targeting spacer (Fig. 4c). The sensitivity of ΦKZ to RNA-targeting but not DNA-targeting underscores the role of the shell in broad spectrum resistance to DNA-cleaving enzymes and also provides the first evidence of a DNA phage being inhibited by Cas13.

Evasion of the endogenous *P. aeruginosa* Type I CRISPR-Cas system by ΦKZ suggests that these jumbo phages are likely to pervasively avoid this system in nature. Supporting this hypothesis, our analysis of >4,000 *P. aeruginosa* non-redundant spacers (Type I-C, I-E, and I-F) reported by van Belkum et al. (2015) found no spacers against ΦKZ, or its jumbo phage relatives ΦPA3, PaBG, KTN4, and PA7 (Table 1). This is in contrast to the many spacer matches from each system against diverse *P. aeruginosa* phages, such as those assayed in our screen and those encoding anti-CRISPR proteins (Table 1). Finally, given the efficacy of the RNA-targeting CRISPR-Cas13 system, we propose that perhaps these CRISPR systems are well-suited to target the mRNA of DNA phages when the DNA is inaccessible (i.e. due to base modifications or physical segregation). However, the rules governing the ability of Cas13 to limit the replication of DNA phages remain to be elucidated as only 3/11 LseCas13a crRNAs tested targeted ΦKZ and 0/6 were effective at targeting phage JBD30 (Extended Data Fig. 3).

**Table 1:** ΦKZ and ΦKZ-like phages have no natural spacers matching their genomes from a natural collection of >4000 *P. aeruginosa* spacers. The total number of Type I CRISPR spacers with a perfect match to the indicated P. aeruginosa phages assayed in this study and previous CRISPR-Cas studies. The experimental sensitivity of each phage to the indicated subtypes are shown. AcrIE3 and AcrIF1 are I-E and I-F anti-CRISPR proteins, respectively. * indicates that all spacers have mismatches (≤4) to the F8 genome.

| Phage | # spacers | Type I CRISPR sensitivity | Reference |
| --- | --- | --- | --- |
| DMS3m | 75 | I-C: sensitive<br>I-E: resistant (AcrIE3)<br>I-F: sensitive | This study, ref. 13,33 |
| JBD30 | 51 | I-C: sensitive<br>I-E: sensitive<br>I-F: resistant (AcrIF1) | This study, ref. 12,13 |
| JBD18 | 51 | I-F: sensitive | Ref. 33 |
| JBD25 | 46 | I-F: sensitive | Ref. 33 |
| JBD68 | 28 | I-C: sensitive | This study |
| D3 | 49 | I-C: sensitive | This study |
| F8 | 3* | I-C: sensitive | This study |
| ΦKZ | 0 | I-C: resistant | This study |
| phiPA3 | 0 | not assayed |  |
| PaBG | 0 | not assayed |  |
| KTN4 | 0 | not assayed |  |
| PA7 | 0 | not assayed |  |

Here, we searched for CRISPR-Cas resistant phages by designing crRNAs against them and testing their efficacy. This screen identified jumbo phage ΦKZ (genome size: 280,334 bp^31^) as resistant to the Type I-C CRISPR system, and subsequently to Type II-A and Type V-A single effector nucleases Cas9 and Cas12. ΦKZ is also recalcitrant to the Type I restriction-modification system of *P. aeruginosa*. Despite this apparent resistance *in vivo,* ΦKZ genomic DNA is sensitive to restriction enzymes and Cas9 cleavage *in vitro*. We propose that the assembly of a proteinaceous compartment to house the replicating phage DNA creates a physical protective barrier resulting in the resistance of phage ΦKZ to DNA-cleaving enzymes (Fig. 4d). Although this shell-like structure has only been documented among the jumbo phages of *Pseudomonas*^28,29^, we consider that physical occlusion of phage genomic DNA through this and other mechanisms may comprise a novel route to immune system evasion in bacteria.

The pan-resistance of ΦKZ to DNA-targeting enzymes provides an explanation for the elaborate and impressive shell structure, and suggests that the phage DNA may never be exposed to the cytoplasm. Other hypotheses to describe the shell’s existence remain to be addressed, including protection from phage-derived nucleases that degrade the bacterial genome or as a mechanism to spatially restrict the large phage genome during replication and packaging. Regardless, we conclude that the nucleus-like shell provides a strong protective barrier to DNA-targeting immune pathways. We expect that simple CRISPR-based screens, such as the one conducted here, may reveal many other fascinating mechanisms that phages have evolved to ensure their replicative success when faced with immune systems to overcome.

## Acknowledgements

Research in the Bondy-Denomy lab was supported by the University of California San Francisco Program for Breakthrough in Biomedical Research, funded in part by the Sandler Foundation, and an NIH Office of the Director Early Independence Award (DP5-OD021344). This work was also supported by HHMI (DAA) and NIH grants R35GM118099 (DAA), and GM104556 (DAA).

ΦKZ, JBD30, JBD68, D3, and F8 were provided by Alan Davidson’s lab. Phage DMS3m was a gift from the O’Toole lab and Jason M. Peters and Carol A. Gross provided the *S. pyogenes cas9* expression plasmid for integration in the PAO1 chromosome. pTE4495 (MbCpf1/MbCas12a) was a gift from Ervin Welker (Addgene plasmid # 80339), and LseCas13a (Addgene plasmid #83486) is a gift from Jennifer Doudna.

## Author Contributions

S.D.M. conducted restriction-modification experiments, constructed Cas13 strains and conducted associated experiments with that strain and the Cas12 strain, including all liquid infection assays, and prepared figures. J.D.B. constructed and conducted experiments with Cas9 and Cas12 expressing strains, and conducted *in vitro* digestion assays. L.M.L. conducted Type I-C Cas3 experiments. J.B.-D. conceived of the project, conducted Cas3 and Cas9 experiments, supervised all experiments, and wrote the manuscript together with S.D.M. E.S.N. conducted microscopy experiments, under the supervision of D.A.A. All authors edited the manuscript.

## Materials & Correspondence

Requests should be made to

## Extended Data Figures

**Extended Data Figure 1:**
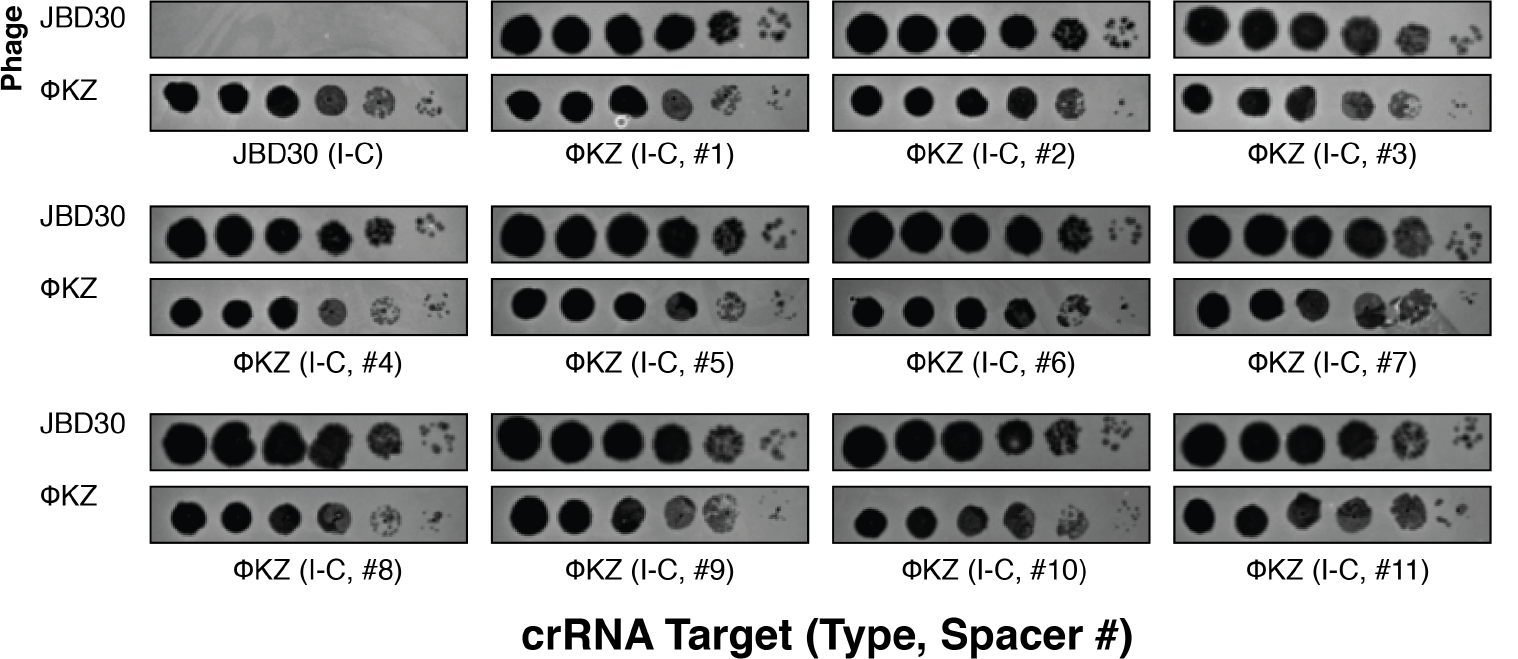
Phage ΦKZ is resistant to Type I-C CRISPR-Cas immunity. *P. aeruginosa* strain PAO1 expressing the Type I-C *cas* genes and crRNAs programmed to recognize the indicated phage. One crRNA targeting JBD30 is shown, while 11 distinct crRNAs are directed towards ΦKZ. Phages are spotted in ten-fold serial dilutions (left to right) on a lawn of the indicated PAO1 strain.

**Extended Data Figure 2:**
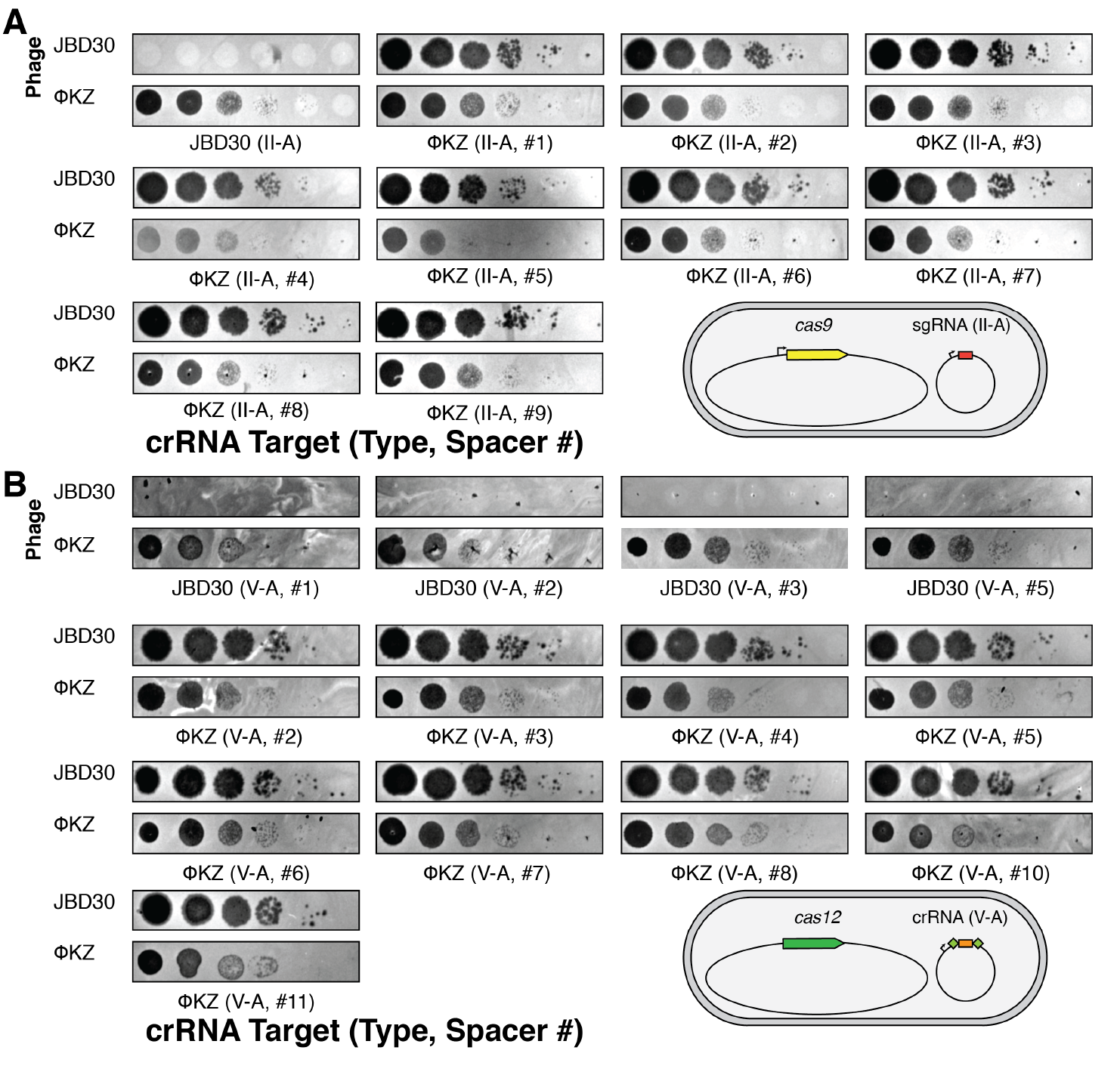
Phage ΦKZ is resistant to Type II-A and V-A CRISPR-Cas immunity. *P. aeruginosa* strain PAO1 expressing the (**a**) Type II-A Cas9 or (**b**) Type V-A Cas12a effectors and sgRNAs/crRNAs programmed to recognize the indicated phage. Phages are spotted in ten-fold serial dilutions (left to right) on a lawn of the indicated PAO1 strain.

**Extended Data Figure 3:**
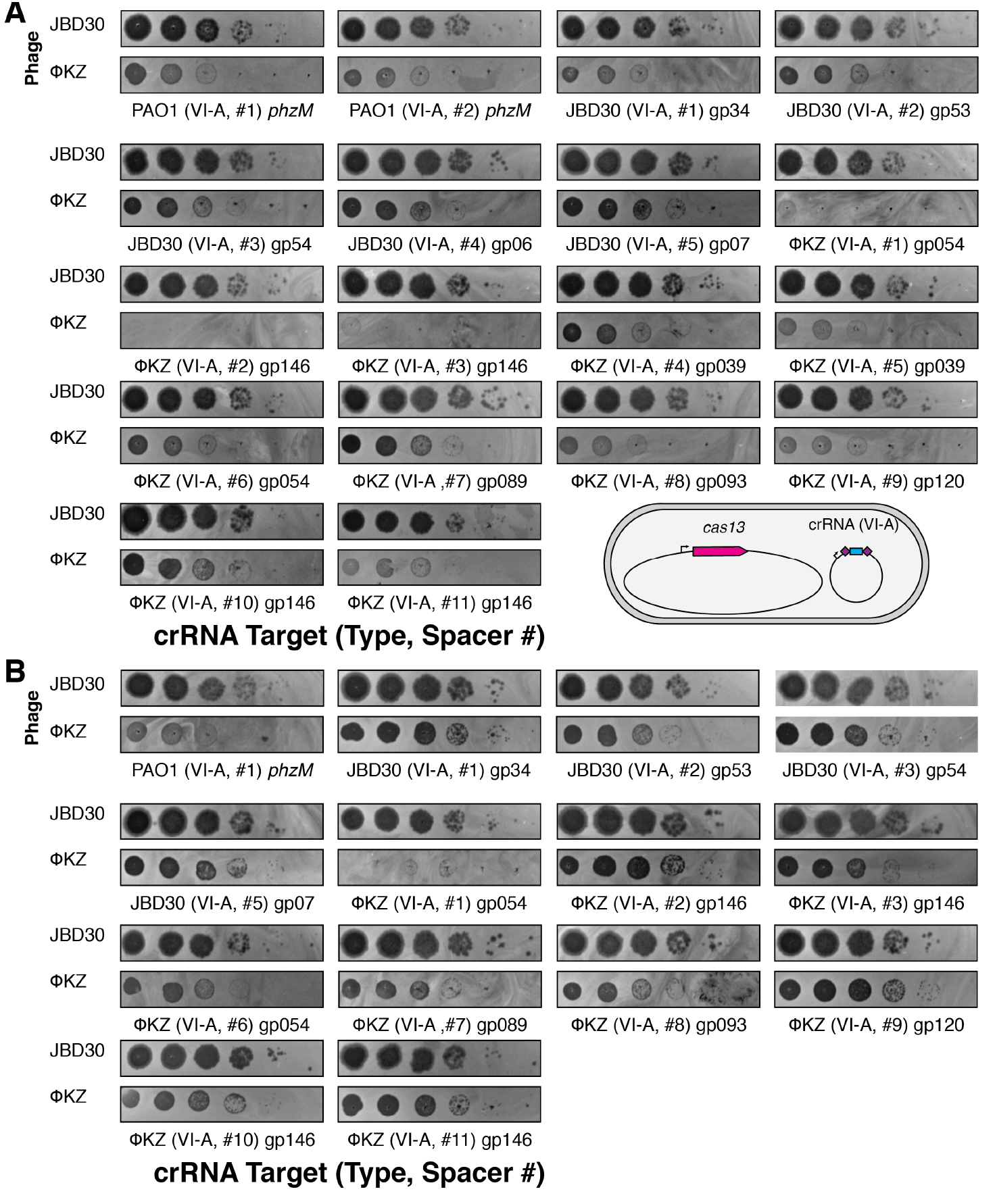
Phage ΦKZ is sensitive to Type VI-A CRISPR-Cas immunity. *P. aeruginosa* strain PAO1 expressing the Type VI-A (a) LseCas13a or (b) LshCas13a effectors and crRNAs programmed to recognize the indicated phage. Phages are spotted in ten-fold serial dilutions (left to right) on a lawn of the indicated PAO1 strain.

## Methods

### Bacterial growth and genetic manipulation

Strains, plasmids, phages, and spacer sequences used in this study are listed in Supplemental Tables 1-4. *Pseudomonas aeruginosa* strain PAO1 was grown in LB at 37°C with aeration at 225 RPM. When necessary plating was performed on LB agar with carbenicillin (200 μg/ml) or gentamicin (50 μg/ml). Gene expression was induced by the addition of L-arabinose (0.1% final) and/or isopropyl β-D-1-thiogalactopyranoside (IPTG, 0.5 mM or 1 mM final). For chromosomal insertions at the attTn7 locus, *P. aeruginosa* cells were electroporated with the integrating vector pUC18T-lac and the transposase expressing helper plasmid pTNS3, and selected on gentamicin. Potential integrants were screened by colony PCR with primers PTn7R and PglmS-down. Electrocompetent cell preparations, transformations, integrations, selections, plasmid curing, and FLP recombinase mediated marker excision with pFLP were performed as described previously^32^.

### Phage growth and DNA extraction

Phage growth was conducted in LB at 37°C with PAO1 as a host. Growth curves were conducted in a Biotek Synergy plate reader at 37°C with orbital shaking set to maximum speed. Phage stocks were diluted and stored in SM buffer^33^ and used for routine plaquing assays. For high titer lysates to generate phage DNA, plates with a high number of plaques were flooded with SM buffer and collected^33^. The lysates were subsequently DNase treated and filtered through a 0.22 μm filter. Phage DNA was extracted with the Wizard Genomic DNA Purification Kit (Promega). DNA restriction assays were performed according to standard NEB protocols and restriction fragments were assessed by agarose gel electrophoresis.

### Type I-C CRISPR-Cas system expression in *P. aeruginosa* PAO1

Type I-C CRISPR-Cas function was tested by electroporating a strain containing I-C *cas* genes with pHERD30T plasmids encoding crRNAs that target phages. To express this system heterologously in PAO1, the four effector *cas* genes (*cas3-5-8-7*) were cloned into pUC18T-lac and inserted in the PAO1 chromosome as described above. After removal of the gentamicin marker, this strain was electroporated with the same pHERD30T crRNA-encoding plasmids to confirm function upon IPTG/arabinose induction.

### crRNA cloning and expression

All crRNAs used here, were cloned into established entry vectors in the pHERD30T background. After removing a pre-existing BsaI cut site in the vector by mutagenesis, a pseudo-CRISPR array (i.e. repeat-spacer-repeat for Type I, V, VI, or a sgRNA scaffold for Type II) was then cloned into the vector, where the spacing sequence possessed two inverted BsaI digest sites, to facilitate scarless cloning of new spacers. Desired spacer sequences were ordered as two complementary oligonucleotides that generate sticky ends when annealed, to be cloned into the entry vector, which was BsaI digested. Spacer oligonucleotides were PNK treated, annealed, and ligated into the entry vector.

### Streptococcus pyogenes (Spy) Cas9 and sgRNA expression in *P. aeruginosa*

The *S. pyogenes Cas9* gene was cloned into a pUC18T-Ara integration vector and then inserted into the attTn7 locus of PAO1. A single guide RNA scaffold was constructed based on a previous design^34^ with internal BsaI cut sites to enable insertion of pre-annealed oligos for scarless sgRNA design. This sgRNA scaffold was amplified with primers p30T-gRNA_BsaI and p30T-gRNA_BsaI_rev. The resulting product was inserted into the pHERD30T vector via Gibson assembly following backbone (pJW1) amplification by inverse PCR with primers gRNA_BsaI-p30T and gRNA_BsaI-p30T-rev. The sgRNA scaffold was positioned into pJW1 so that following BsaI cleavage the spacer insert +1 position would conincide with the pBAD TSS +1 position. The resulting plasmid, pJB1, was BsaI digested (NEB) followed by ligation of indicated pre-annealed oligos. Table 3 contains a complete list of all target sequences. The sequence of the sgRNA construct with BsaI site locations is shown in Supplemental Table 3.

### Cas9 in vitro cleavage

Cas9-based phage genome cleavage *in vitro* was conducted with purified Cas9 protein (NEB #M0386S), and the Cas9-gRNA-tracrRNA based cleavage reaction was then performed using according to the manufacturer’s (NEB) instructions. Cas9 crRNAs (Supplemental Table 3) were ordered as Alt-R CRISPR-Cas9 crRNAs from IDT and utilized without further modification. The tracrRNA was amplified using primers tracrRNA-FOR and tracrRNA-REV from a plasmid (pBR62). The tracrRNA was produced through a T7 RNAP reactions using dsDNA encoding the tracrRNA downstream of a T7 RNAP promoter. Cas9 protein (NEB) was combined with pre-annealed crRNA and tracRNA complex at a 1:1 molar ratio. The reaction was performed at 37°C for 4 hrs with 300 ng of ΦKZ or DMS3 genomic DNA and the products were assessed by agarose gel electrophoresis. Two Cas9 guides were selected that would cleave at pos. 158,649 and 168,715 of the ΦKZ genome to liberate an ~10 kb fragment.

### Cas12a and crRNA design for PA expression

The humanized allele of the *cpf1* gene of *Moraxella bovoculli* (MBO_03467, KDN25524.1) was sub-cloned from pTE4495 (Addgene) into pUC18T-lac using primers pUC_cpf1_F and pUC_cpf1_R and inserted in the PAO1 chromosome as described above. A Cpf1 repeat-spacer-repeat pseudo-CRISPR array was synthesized as oligonucleotides, annealed, and ligated into a pHERD30T vector, digested with NcoI and HindIII. Spacer sequences were cloned into the resulting vector (pJB2) following BsaI digestion and ligation of pre-annealed spacer oligonucleotide pairs.

### Cas13a and crRNA design for PA expression

The wild type allele of the *cas13* gene of *Listeria seeligeri* and *Leptotrichia shahii* were sub-cloned from p2CT-His-MBP-Lse_C2c2_WT and p2CT-His-MBP-Lsh_C2c2_WT (Addgene) into pUC18T-lac. LseCas13 and Lsh Cas13 were inserted in the PAO1 chromosome as described above. An Lse and an Lsh Cas13a repeat-spacer-repeat pseudo-CRISPR array were synthesized as oligonucleotides, annealed, and ligated into a pHERD30T vector, digested with NcoI and EcoRI. Spacer sequences were cloned into the resulting vectors (pSDM057 and pSDM070, respectively) following BsaI digestion and ligation of pre-annealed spacer oligonucleotide pairs. crRNA expression vectors were introduced into PAO1 tn7::*cas13*^Lse^ and PAO1 tn7::*cas13*^Lsh^. The resulting strains were grown to saturation in LB at 37 °C. 4 mL of 0.7% agar, 10 mM MgSO_4_, 1 mM IPTG molten top agar were seeded with 100 μL saturated culture and spread on 20 mL 10 mM MgSO_4_, 50 μg/mL gentamicin, 0.1% (L)-arabinose, 1 mM IPTG LB agar plates. 2.5 μL 10-fold serial dilutions of bacteriophage JBD30 and ΦKZ were spotted on plates. Plates were incubated at 37 °C overnight and were imaged on the following day.

### Restriction-Modification Assay

The PAO1 *hsdR* gene (PA2732) was knocked out using CRISPR-Cas9 and a targeted sgRNA. PAO1, PAK, and PAO1 Δ*hsdR* were grown to saturation in LB at 37 °C. 4 mL of 0.7% agar, 10 mM MgSO_4_ molten top agar were seeded with 100 μL saturated culture and spread on 20 mL 10 mM MgSO_4_ LB agar plates. 2.5 μL 10-fold serial dilutions of bacteriophage JBD30 and ΦKZ propagated on strain PAO1 and PAK were spotted on plates. Plates were incubated at 37°C overnight and were imaged the following day.

### Immunofluorescence

#### Sample Growth

5 mL overnight cultures of a strain expressing Cas9 and an sgRNA targeting ΦKZ (SDM065) and a strain expressing cMyc-ORF152 (bESN27) were grown at 30 °C in LB media with gentamicin. A 1:30 back-dilution of the overnight culture into LB was grown at 30°C for 1 h. Protein and guide expression was induced with 0.1% arabinose and 0.5 mM IPTG, respectively. After 1 h of expression, an aliquot of uninfected cells was fixed while the remaining cultures were infected with ΦKZ using MOI 1.5. Infected cell aliquots were collected and fixed at 60 mpi.

#### Fixation

This protocol was adapted from ref. 35. Samples were fixed with 5X Fix Solution (12.5% paraformaldehyde, 150 mM KPO4, pH 7.2) and incubating for 15 minutes at room temperature followed by 20 minutes on ice. Samples were then washed in PBS 3 times and finally resuspended in GTE (50 mM glucose, 10 mM EDTA, pH 8.0, 20 mM Tris-HCl, pH 7.65) with 10 ug/mL lysozyme. Resuspended cells were transferred to polylysinated coverslips and dried. Once dry, coverslips were incubated in cold methanol for 5 minutes followed by cold acetone for 5 minutes. Cells were rehydrated by a rinse in PBS followed by a 3-minute incubation in PBS + 2% BSA blocking solution. Cells were incubated with a 1:50 dilution of primary antibody (Cas9 (7A9-3A3): sc-517386 or cMyc (9E10): sc-40) in PBS + 2% BSA for 1 hr followed by 3, 7 minute washes in fresh PBS + 2% BSA. Coverslips were then incubated in the dark for 1 hr with secondary antibody (goat anti-mouse Alexa Fluor 555, Life Technologies A-21424) diluted 1:500 in PBS + 2% BSA. DAPI was added for the final 10 minutes of the incubation. Cells were washed in PBS 3 times for 7 minutes. Coverslips were then placed on slides using mounting media (v/v 90% glycerol, v/v 10%Tris pH 8.0 and w/v 0.5% propyl-gallate) and sealed with clear nail polish.

#### Microscopy and Analysis

Images were collected using a Zeiss Axiovert 200M microscope.

## Supplementary Information

### Strains, Plasmids, and Oligonucleotides

**Supplemental Table 1.** Phage and Strains.

| Name | Source | Reference |
| --- | --- | --- |
| ΦKZ | Davidson Lab | 31 |
| DMS3m | O'Toole Lab | 33 |
| JBD30 | Davidson Lab | 36 |
| F8 | Davidson Lab | Unpublished<br>Accession: DQ163917 |
| D3 | Davidson Lab | 37 |
| JBD68 | Davidson Lab | 36 |
|  | Genotype | Reference |
| JW31 | PAO1 tn7::I-C/Cas3<sup>PA</sup> | This study |
| JB10 | PAO1 tn7::cas9<sup>Spy</sup> | This study |
| JB90 | PAO1 tn7::cas12a<sup>Mb</sup> | This study |
| SDM084 | PAO1 tn7::cas13<sup>Lse</sup> | This study |
| SDM020 | PAO1 ΔhsdR | This study |

**Supplemental Table 2.** Plasmids.

| Name | Information | Ref. |
| --- | --- | --- |
| pHERD30T | Arabinose inducible, gentamicin resistant shuttle vector | 38 |
| pUC18T-Lac | IPTG inducible, Amp<sup>R</sup>/Gent<sup>R</sup>, Tn7 integrative plasmid with FRT sites flanking gentamicin cassette | 39 |
| pTNS3 | Expresses tnsABCD for flipping out cassette flanked by FRT sites | 40 |
| pBR62 | Contains Spy Cas9 tracrRNA sequence | This study |
| pJW1 | pHERD30T with BsaI site at pos. 235 GAGACC mutated to GTGACC | This study |
| pJW13 | pJW1 with Type I-C pseudo-CRISPR array for spacer cloning | This study |
| pJB1 | pJW1 with Type II-A sgRNA backbone at the +1 TSS of pBAD | This study |
| pJB2 | pJW1 with Type V-A pseudo-CRISPR array for spacer cloning | This study |
| pSDM057 | pJW1 with Type VI-A (LseCas13a) pseudo-CRISPR array for spacer cloning | This study |
| pSDM070 | pJW1 with Type VI-A (LshCas13a) pseudo-CRISPR array for spacer cloning | This study |

**Supplemental Table 3.**
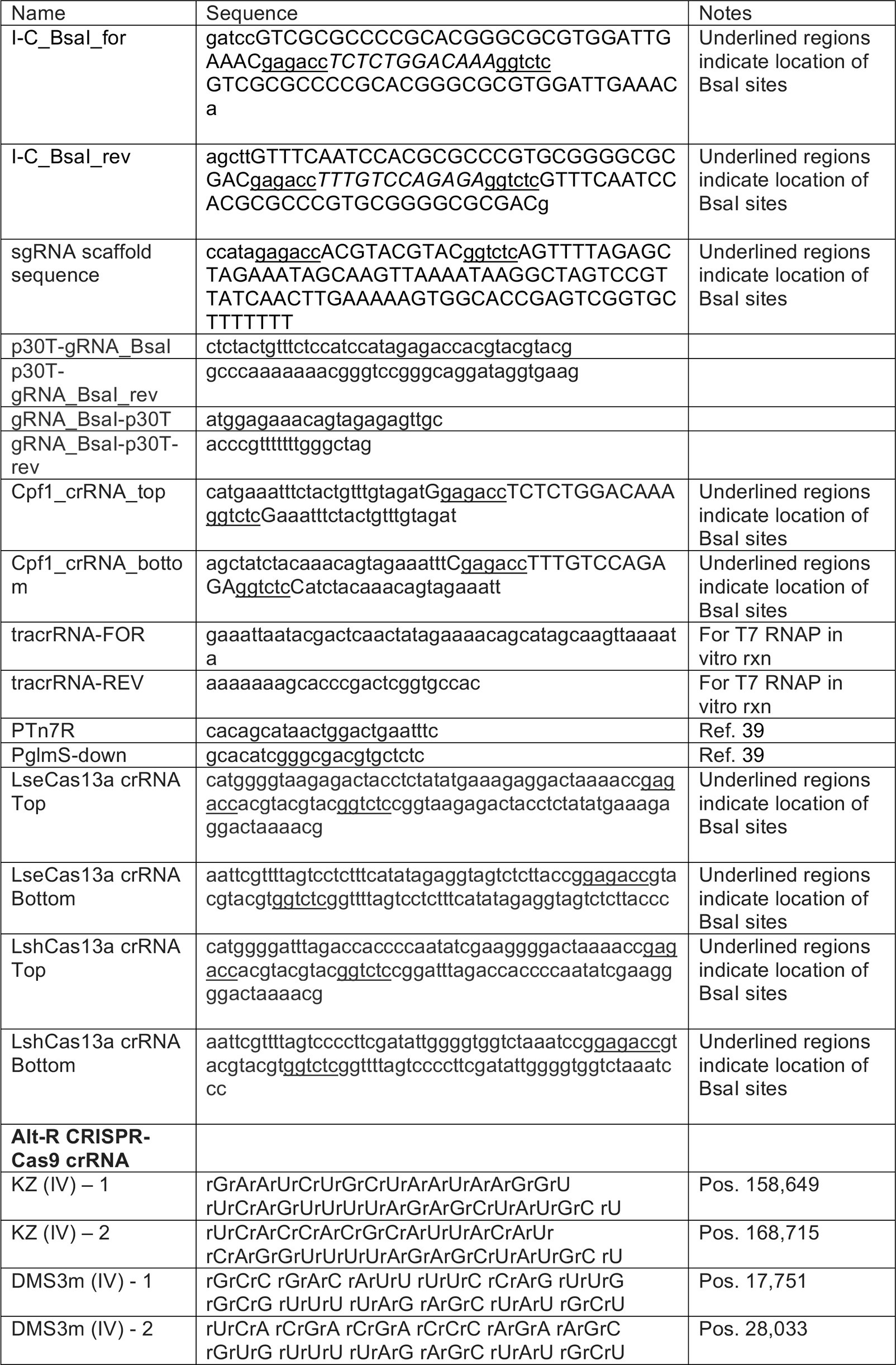
Oligonucleotides, g-blocks, crRNAs.

**Supplemental Table 4.**
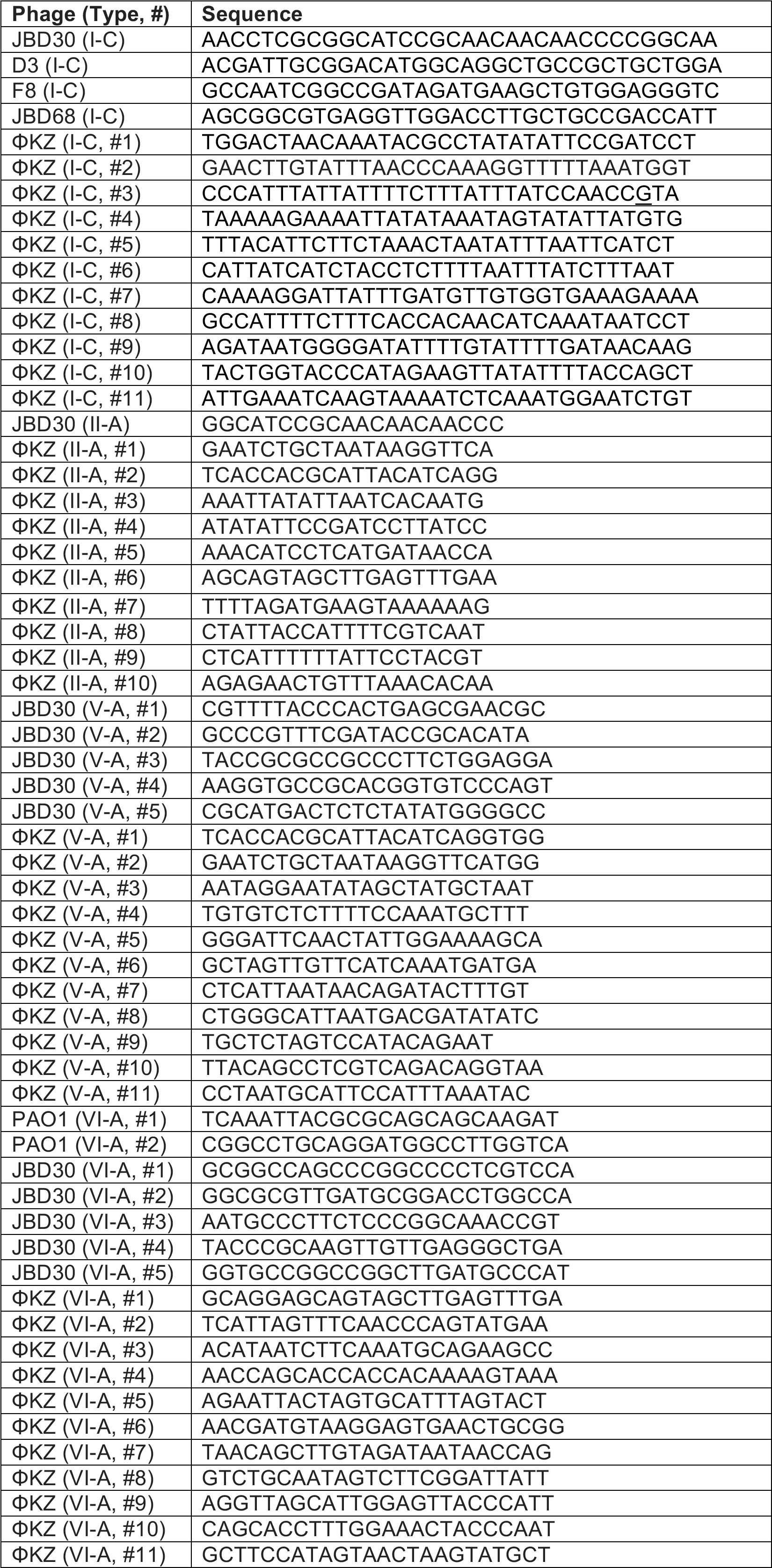
Spacer sequences.

